# Detrital Food Web Drives Aquatic Ecosystem Productivity in a Managed Floodplain

**DOI:** 10.1101/610055

**Authors:** Carson A. Jeffres, Eric J. Holmes, Ted R. Sommer, Jacob V.E. Katz

## Abstract

Differences in basal carbon sources, invertebrate density and salmon growth rate were observed in food webs across a lateral transect of aquatic habitats in the Sacramento River Valley, California. Similar to many large river valleys globally, the Sacramento River Valley has been extensively drained and leveed, hydrologically divorcing most floodplain wetlands and off-channel aquatic habitats from river channels. Today, the former floodplain is extensively managed for agriculture and wildlife habitat. Food web structure and juvenile Chinook Salmon (O*ncorhynchus tshawytscha*) growth were compared in three aquatic habitat types–river channel, a perennial drainage canal in the floodplain, and agricultural floodplain wetlands, which was seasonally inundated to provide bird and fish habitat during the non-agricultural growth season (late winter). Zooplankton densities on the floodplain wetland were 53 times more abundant, on average, than in the river. Juvenile Chinook Salmon raised on the floodplain wetland grew at mm/day, a rate 5x faster than fish raised in the adjacent river habitat (0.18 mm/day). Mean water residence times calculated for the floodplain agricultural wetland, perennial drainage canal and Sacramento River were 2.15 days, 23.5 seconds, and 1.7 seconds, respectively. Carbon in the floodplain wetland food web was sourced primarily through heterotrophic detrital pathways while carbon in the river was primarily autotrophic and sourced from in situ phytoplankton production. Hydrologic conditions typifying the ephemeral floodplain-shallower depths, warmer water, longer residence times and detrital carbon sources compared to deeper, colder, swifter water and an algal-based carbon source in the adjacent river channel-appear to facilitate the dramatically higher rates of food web production observed in floodplain verses river channel habitats. These results suggest that hydrologic patterns associated with winter flooding provide Mediterranean river systems access to detrital carbon sources that appear to be important energy sources for the production of fisheries and other aquatic resources.

## Introduction

The benefits of annual floodplain inundation to riverine ecosystems and fish populations are well recognized in relatively unaltered tropical river systems [1, 2]. Floodplains and other seasonally-inundated off-channel river habitats have not been as thoroughly studied in temperate climates [3]. In Europe and North America, levees have been constructed along almost every major lowland river to allow development of fertile floodplains for farms and cities [4]. In the Central Valley of California approximately 95% of the historic floodplain wetlands have been drained or are no longer accessible to aquatic species behind ~3360 km of state and federal levees [5]. This landscape-scale hydrologic divorce of river channel and floodplain has only recently been widely recognized and ecosystem responses have only just begun to be studied and quantified [6]. Stream metabolism is a means of measuring how energy is created and used within aquatic ecosystems. Here we employ stream metabolism techniques to compare and contrast how energy (carbon) flows through aquatic food webs in three aquatic habitat types in the Sacramento River Valley in California, USA (Fig 1) typical of those found in leveed river valleys globally.

The Sacramento Valley is a Mediterranean climate where summers are long and dry and almost all precipitation falls in winter and spring. This results in rivers with high annual and seasonal variability in flows with flooding occurring exclusively during the winter/spring wet season. Autotrophic production is widely recognized as an important driver of aquatic food web productivity. Water temperatures on Central Valley floodplains inundated during winter flood season are generally warmer due to decreases in depth, and increases in surface area and water residence time compared to the relatively deep, cool and swift river channel [7, 8]. Floodplain habitats provide greater food resources for grazing zooplankton, which ultimately provide food resources for fishes [7–10]. A significant portion of the trophic energy transfer in floodplains and similar off-channel habitats may move through heterotrophic food webs driven by breakdown of plant detritus [11–15]. Studies on the Yolo Bypass in California have found that detritivores such as chironomid larvae can be very abundant in hydrologically-activated floodplain habitats during flood events [16]. The abundant floodplain food resources contribute significantly to the diets of juvenile salmonids accessing these off-channel habitats during flood events [16, 17] where they grow more quickly than fish confined to adjacent leveed river channels [10, 18].

This study identifies differences in hydrology, carbon source and productive capacity of food webs in three aquatic habitats in the Sacramento River Valley (Fig 1) that epitomize a typical lateral cross section of a developed agricultural river valley: seasonally inundated agricultural floodplain wetlands, perennial canals engineered to drain the floodplain surface, and leveed river channels. We hypothesized that access to terrestrial detrital carbon sources would contribute to higher rates of heterotrophic food web productivity in ephemerally inundated floodplain wetland habitats compared to the adjacent perennial drainage canal or river channel habitats. Higher rates of ecosystem productivity should lead us to observe an aquatic food web characterized by:

- Higher densities of zooplankton on the floodplain compared to the adjacent perennial canal or river channel habitats;
- Increased growth rates of juvenile salmon reared on the floodplain habitat compared to the perennial canal or river channel habitats;

## Methods

### Study Area

The Sacramento River is the largest river in California draining 71,432 km^2^. Seasonal flow events from winter rains and spring snow-melt historically inundated much of the Sacramento Valley floodplain. Starting in the late 1800s levees were constructed to protect agricultural and urban development. Today, less than 5% of historical floodplain wetlands remain [19]. The exception to this hydrological disconnection of the Sacramento River from its floodplain is a series of landscape-scale flood protection projects designed to bypass flood waters around critical urban and agricultural areas. By diverting high flows on to specifically managed floodways these “bypasses” alleviate flood stress on critically important downstream levees. The largest bypass is the Yolo Bypass, a 24,000 ha floodway adjacent to the city of Sacramento and immediately upstream of the Sacramento-San Joaquin Delta (Fig 1). Yolo Bypass is extensively farmed during summer and floods in two out of three winters, on average, although flood events are frequently shorter than a week [17]. There are also substantial areas of the Yolo Bypass that are managed as seasonal agricultural floodplain wetlands to support wildlife, as well as a perennial drainage canal that spans much of the eastern edge of the floodplain.

**Fig 1.**
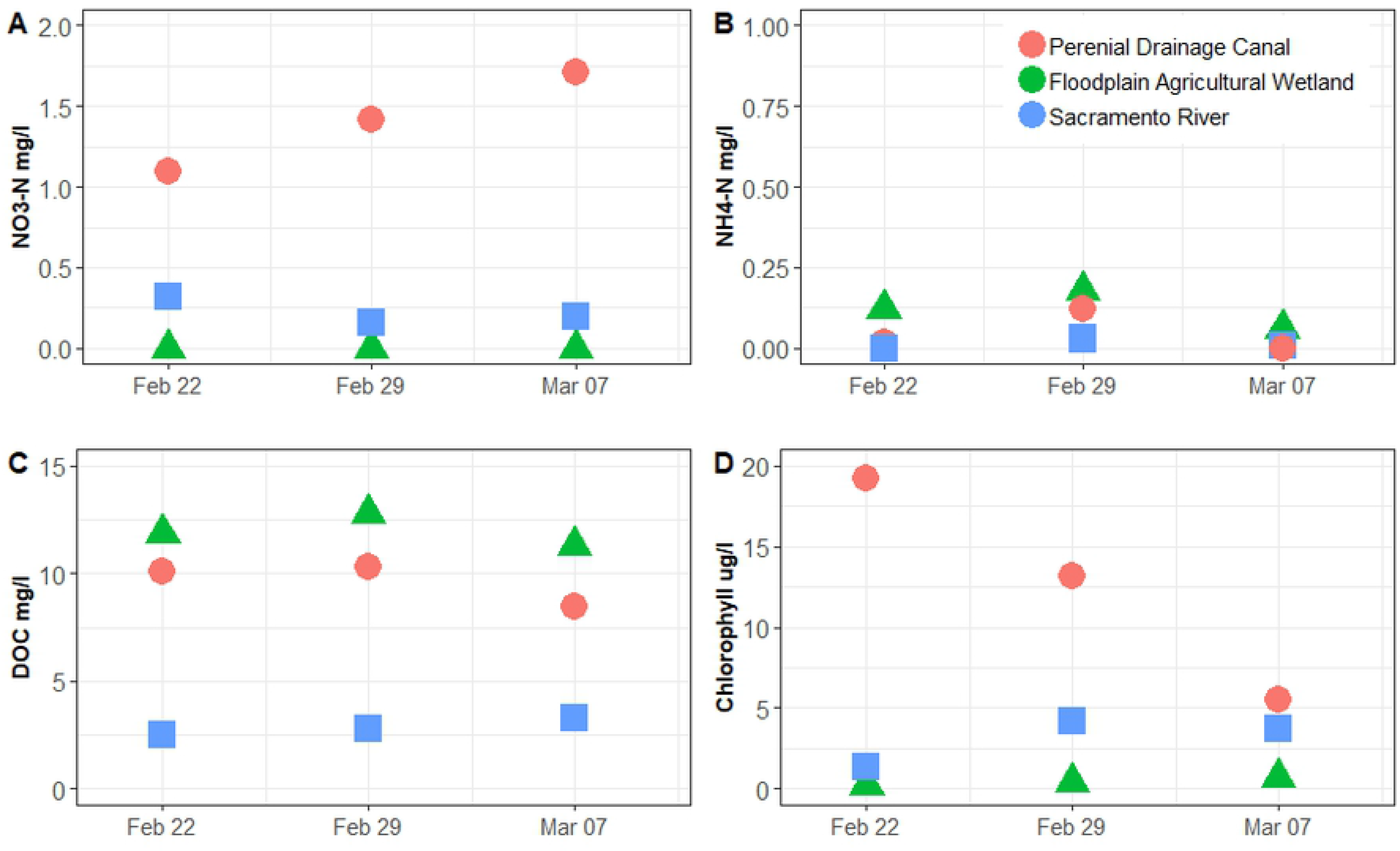
Location of study sites (blue stars) from left to right – flooded floodplain agricultural wetlands, perennial channel, and Sacramento River.

### Hydrology and water chemistry

Hydrology and water chemistry data were collected in order to document physical habitat differences between each of the sites, and to provide the basis for stream metabolism modeling (below). Hourly discharge and stage data in the Sacramento River and Yolo Bypass perennial drainage canal were downloaded from the California Data Exchange Center (cdec.ca.gov) at stations “VON” and “YBY”, respectively. Water residence times in the Sacramento River and the Yolo Bypass perennial drainage canal sites were calculated using a relationship between cross-sectional area and mean velocities at a variety of flows at the stations. Then that relationship was used to calculate mean water velocities from the flows that were observed during this study. Data was collected from published values at usgs.gov for site numbers 11425500 and 11453000, respectively. Water quality data were collected via a handheld YSI EXO sonde that measures water temperature, dissolved oxygen, salinity, pH, conductivity, and turbidity. In addition, at each location an Onset Hobo U26 dissolved oxygen and temperature data logger (Onset Computer Corporation, Bourne, MA) was placed near the cages which collected data at 15-minute intervals throughout the study. Water column chlorophyll samples were collected in 1-liter HDPE bottles and brought back on ice to the laboratory for analysis. Chlorophyll samples were filtered upon return to the laboratory and the filters were frozen until further analysis. Water column nutrient samples were collected in 125 ml HDPE bottles, placed on ice, and then refrigerated when returned to the laboratory; constituents measured were total phosphorous, phosphate-phosphorous, ammonia-nitrogen, nitrate-nitrogen, total nitrogen, and dissolved organic carbon. Dissolved constituents were determined on samples following filtering through a 0.2 μm Millipore polycarbonate membrane filter; total concentrations were determined on non-filtered samples.

### Food web

Experimental enclosures were used as a tool to compare the food web responses across the three habitats. These enclosures were constructed with a rigid frame of 25.4 mm pvc pipe that was wrapped with an extruded plastic mesh with openings of 6.3 mm. The 6.3 mm mesh allowed for free movement of zooplankton and other invertebrates while keeping fish inside of enclosures. The plastic mesh was secured to the pvc frame with plastic zip ties. Enclosures were 1.2 m long × 1.2 m wide × 0.6 m deep. Enclosures were placed into each of the three study locations (agricultural floodplain wetland, perennial floodplain channel, and Sacramento River; Fig1). Enclosures in the agricultural floodplain wetlands, where water depth did not fluctuate, were secured to metal posts driven into the ground with their top mesh even with the water surface. Floats were attached to the top of the cages in the canal and Sacramento River causing the cages to float with the top mesh directly at the water surface. These floating enclosures were attached to wood pilings via a tethered line that allowed the cages to float at the water’s surface as water elevations changed. The agricultural floodplain wetland location consisted of a 0.81 ha farm field that had a single inflow from a supply ditch and a single exit. Water depth was maintained at approximately 0.46 m by boards placed into an irrigation box for a total standing water volume of approximately 3722 m^3^ in the field. The field, usually planted to rice, had been fallow the previous year when naturally recruited herbaceous vegetation had been allowed to grow. Previous studies have shown similar zooplankton composition in fallow fields and post-harvested rice fields at the study location [20]. The field was flooded using water from the adjacent irrigation canal on January 18, 2016, 32 days prior to the start of the project.

Juvenile Chinook Salmon from the Feather River hatchery were transported to the study site via a large fish transporting tank where they were tagged with 8 mm passive integrated transponder (PIT) tags implanted into the abdominal cavity [21]. This study was carried out in strict accordance with the recommendations in the UC Davis’s Institutional Animal Care and Use Committee (Protocol # 18883). At the end of the experiment, fish were euthanized with a quick blow to the head as approved in our IACUC protocol and placed on ice and frozen for further tissue analysis. Fish were held for two days in large plastic tanks at the agricultural wetland site to ensure fish health and successful tag retention prior to planting into the enclosures. Fish were scanned for PIT tag ID, measured, weighed, and placed into enclosures on February 19, 2016. Three enclosures were placed at each site, and 10 fish were placed into each enclosure. Fish were then sampled on days 9, 16, and 23 after initial planting. During sub-sampling, fish were removed from the enclosure, placed into a cooler, individually scanned, measured, weighed, and then placed back into the enclosure. Following the final measurement on March 11, 2016, fish were euthanized and immediately frozen for future gut content and stable isotope analysis for a concurrent project.

Zooplankton and macro-invertebrate samples were collected weekly at each site. To collect zooplankton, a 30 cm diameter 153 μm mesh net was used [22, 23]. The net was attached to a 5 m rope that was thrown into the water and retrieved while maintaining the entire net below the water surface. The net was thrown four times and all four throws were composited into a single sample. Following the fourth retrieval, the sample was rinsed from the net into the collecting cup that was then rinsed into a Whirl-Pak bag, preserved with 95% ethanol and stained with rose bengal. Samples were then taken back to the laboratory for enumeration and identification. Samples were rinsed through a 150 μm mesh and then emptied into a beaker. The beaker was then filled to a known volume to dilute the sample, depending on the density of individuals within the sample, and then sub-sampled with a 1 ml large bore pipette. If densities were still too great for enumeration the sample was split using a Folsom splitter before sub-sampling with the pipette. The dilution volume, number of splits, and number of aliquots removed were recorded and used to obtain total estimates of invertebrates. Zooplankton samples were sorted until a minimum of 500 individuals were counted within a complete sub-sample. If less than 500 individuals were counted, another subsample was enumerated. Invertebrates were identified with the aid of a dissecting microscope at 4-times magnification to the lowest taxonomic level possible using keys [24–26]. Copepods were only identified to family. Terrestrial invertebrates were rare and not included in final counts.

### Stream Metabolism Modeling

Aquatic ecosystem metabolism is often expressed as net ecosystem productivity (NEP), which is calculated as the difference between all photosynthetic energy produced in the system (gross primary productivity, GPP) and the sum of all energy used by organisms (ecosystem respiration, ER). To determine basal carbon sources and identify pathways of carbon transfer through the food web, we calculated biomass and production rates of primary producers (phytoplankton) as well as biomass and consumption rates of primary consumers (zooplankton). In addition to mass balance flux calculations, we used oxygen fluctuations to model stream metabolism and estimate autotrophic and heterotrophic metabolism, which can be used as a proxy to quantify different carbon sources.

Primary production biomass (PB) was calculated using an empirical model where C: Chlorophyll-a was calculated as (32 mg C mg ^−1^ Chlorophyll-a) [27] and production equation as described in Lopez, Cloern (28) (Table 1). Zooplankton biomass (ZB) was calculated by obtaining dry weight from literature values. If the exact species could not be found, an average dry weight of the published genera was used for the calculation. To determine the amount of carbon for each zooplankton taxa, a constant of 0.48 times dry weight was used as described in Andersen and Hessen (29). Phytoplankton primary productivity (PP) and zooplankton grazing rate (ZG) was calculated using equations in Lopez, Cloern (28). Daily surface irradiance data were obtained from the nearby Davis station (station 6) at http://www.cimis.water.ca.gov. Daily surface irradiance was averaged throughout the study period at 138 W/m^2^ and then converted to Einsteins (E) for calculations. Because no in situ measurements were collected for light attenuation within the water bodies, an attenuation coefficient of 1 was used for all calculations for all three habitats as an approximation throughout the study. This value was chosen because of the variability in turbidity due to variations in flow and wind throughout the study and represents an average value compared to other values in the literature.

**Table 1.** Computation used for biomass, production and carbon flow. Computations and table adapted from Lopez, Cloern (28).

| Index | Description | Units | Computation |
| --- | --- | --- | --- |
| PB | Plankton Biomass | mg C m<sup>-3</sup> | = 32(Chl a) |
| ZB | Zooplankton Biomass | mg C m<sup>-3</sup> | = $\sum f_z DW_i / 1000$ |
| PP | Phytoplankton Primary Productivity | mg C m<sup>-3</sup>d<sup>-1</sup> | = (1/H) (0.85 Pg – 0.015 PB*H);<br>Pg = 3.36(Chl a) (E/k) |
| ZG | Zooplankton Grazing Rate | mg C m<sup>-3</sup>d<sup>-1</sup> | = $0.95 m_i^{0.8} e^{\alpha(T-T')} (1 - e^{-0.01PB})$ |
| ZB:PB | Potential Grazing Pressure | fraction | = ZB/PB |
| ZG:PB | Phytoplankton Biomass Grazed Daily | fraction d<sup>-1</sup> | = ZG/PB |
| ZG:ZB | Zooplankton Daily Ration | fraction d<sup>-1</sup> | = ZG/ZB |
| ZG:PP | Primary Production Grazed Daily | fraction | = ZG/PP |

Chl a, chlorophyll a (mg m<sup>-3</sup>); ai, abundance of zooplankton taxon i (number m<sup>-3</sup>); mi, carbon biomass (μg) of zooplankton taxon i; fz, 0.48 from Andersen and Hessen (29); DWi, dry weight (μg) of zooplankton taxon i; Pg, areal gross primary productivity; H, mean water depth (m); E, daily surface irradiance (Einsteins m<sup>-2</sup> d<sup>-1</sup>, PAR); k, attenuation coefficient (m<sup>-1</sup>); T, water temperature (°C); T', 10°C (or 16°C for rotifers).

Stream metabolism was modeled using the StreamMetabolizer package (version 0.10.8) in R (version 3.3.2) [30]. Data manipulation included using a spline interpolation to transform hourly surface irradiance data at http://www.cimis.water.ca.gov to 15-minute data. All interpolated values that were less than zero were replaced with a zero value. The Bayesian model “bayes” within StreamMetabolizer was used and inputs included dissolved oxygen (mg/L), dissolved oxygen (percent saturation), depth (m), water temperature (degrees C), surface irradiance (μmol/m^2^/sec), and discharge (m^3^/sec) from sources described previously.

### Statistical analysis

For analysis of fish size at initial planting, Shapiro Wilks test showed that the data were not normally distributed, so outliers were removed and a non-parametric Kruskal-Wallis test was used to determine variation between treatment groups at the start of the experiments. To determine differences in growth rates between habitats over time, statistical analysis was conducted in JMP Pro version 12.0.1. Because the fish were individually marked and measured several times throughout the experiment, a mixed model repeated measures analysis was used to determine differences in growth between study habitats. Time and location were the fixed effects in the model used. Only fish that were sampled throughout the study were included into the analysis.

## Results

### Hydrology

There was no measurable velocity across the floodplain agricultural wetland, and discharges into and out of the field ranged from zero to 0.02 m^3^s^−1^ throughout the study. Discharge in the floodplain perennial drainage canal ranged from 0.82 m^3^s^−1^ to 68.0 m^3^s^−1^, while discharge in the Sacramento River ranged from 314.3 m^3^s^−1^ to 1,427.2 m^3^s^−1^. Stage in the floodplain agricultural wetland was stable throughout the study, while stage varied by 2.75 m in the canal and 5.00 m in the Sacramento River. Mean residence times calculated for the floodplain agricultural wetland, perennial drainage canal and Sacramento River were 2.15 days, 23.5 seconds, and 1.7 seconds, respectively. Water temperature was most variable on the floodplain agricultural wetland habitat with the highest highs and the lowest lows, and the Sacramento River habitat was the most stable with little fluctuation throughout the study (Fig 2, Table 2). Similar to temperature, dissolved oxygen was highly variable in the floodplain agricultural wetland habitat and very stable in the Sacramento River (Fig 3). Dissolved oxygen in the floodplain agricultural wetland was below saturation throughout most of the study with the exception of days when wind mixed the shallow water. Dissolved oxygen in the perennial drainage canal had a similar pattern to that of the floodplain agricultural wetland, but with smaller daily fluctuations and a higher average dissolved oxygen concentration.

**Fig 2.**
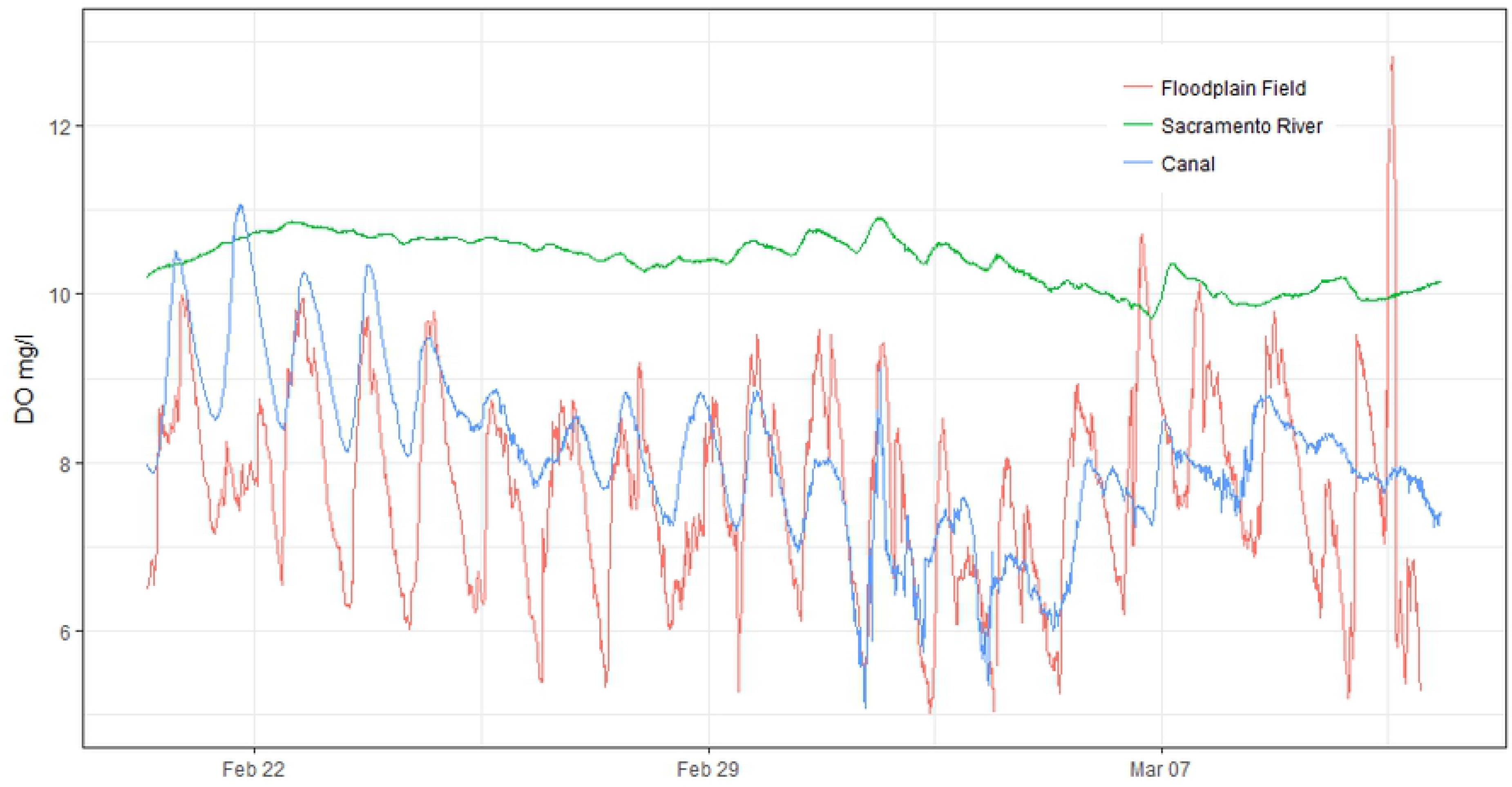
Water Temperature (degrees Celsius) continuously sampled every 15 minutes at flooded floodplain agricultural wetland, perennial drainage canal and Sacramento River habitats.

**Fig 3.**
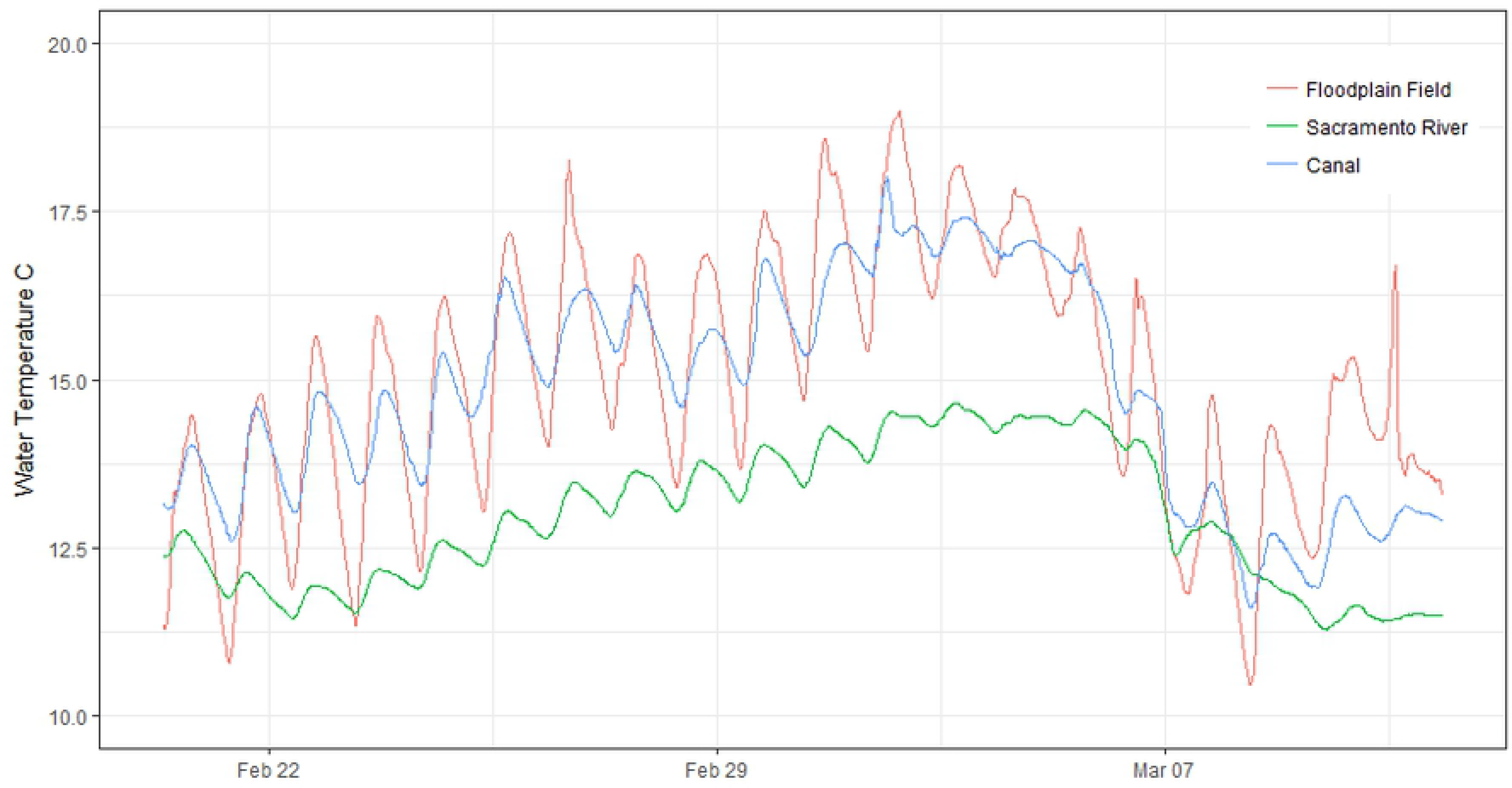
Dissolved Oxygen (DO) in mg/l continuously sampled every 15 minutes at flooded floodplain agricultural wetland, perennial drainage canal and Sacramento River habitats.

**Table 2.** Mean values of water quality data collected from the floodplain agricultural wetland, perennial channel, and Sacramento River study sites.

| Water Quality Constituent | Floodplain agricultural wetland | Perennial channel | Sacramento River |
| --- | --- | --- | --- |
| Average Temperature (°C) | 15.22 | 15.28 | 13.21 |
| Maximum Temperature (°C) | 19.00 | 18.02 | 14.64 |
| Minimum Temperature (°C) | 10.78 | 12.60 | 11.44 |
| Average Dissolved Oxygen (mg/l) | 7.67 | 8.11 | 10.46 |
| **Maximum Dissolved Oxygen (mg/l)** | 10.71 | 11.07 | 10.91 |
| **Minimum Dissolved Oxygen (mg/l)** | 4.86 | 5.07 | 9.72 |
| **Average Total Phosphorous (mg/l)** | 0.21 | 0.90 | 0.11 |
| **Average Phosphate-Phosphorus (mg/l)** | 0.14 | 0.29 | 0.02 |

### Water chemistry

Nitrate-nitrogen values were highest in the perennial drainage canal throughout the study and ranged from 1.10 to 1.71 mg/l (Fig 4). Sacramento River nitrate-nitrogen values were relatively stable and ranged from 0.16 to 0.32 mg/l. Nitrate-nitrogen values were non-detectible (<0.01 mg/l) for all samples collected in the floodplain agricultural wetland throughout the study. Ammonium values were relatively low across all of the habitats sampled, with the highest values being in the floodplain agricultural wetland habitat, ranging from 0.06 to 0.12 mg/l (Fig 4). Dissolved organic carbon (DOC) was highest in the floodplain agricultural wetlands, which ranged from 11.3 to 12.8 mg/l (Fig 4). The Sacramento River had the lowest DOC values, which ranged from 2.5 to 3.3 mg/l.

**Fig 4.**
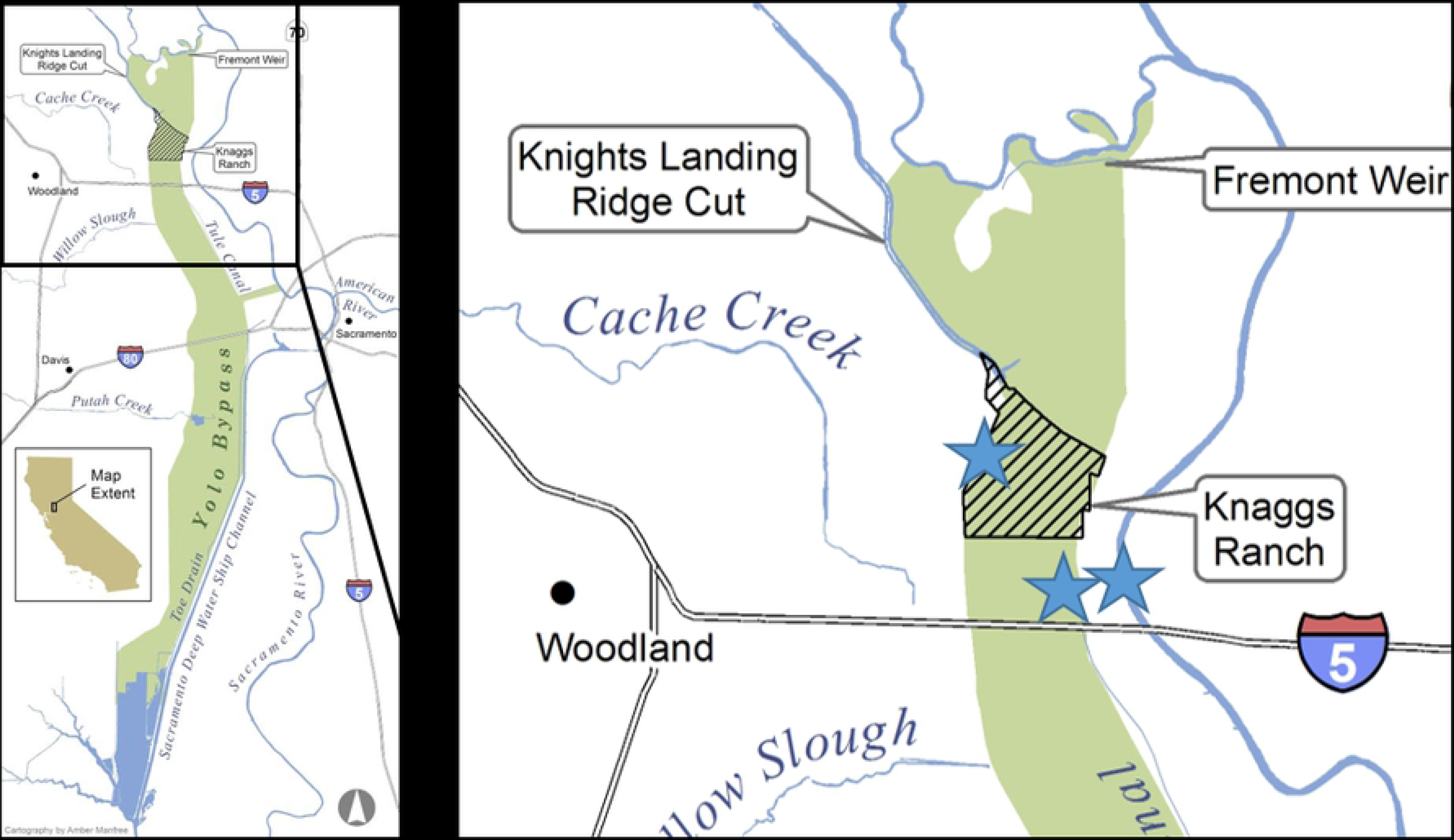
Nitrate-nitrogen (NO_3_-N) mg/l (A), Ammonium-nitrogen (NH_4_-N) mg/l (B), Dissolved organic carbon (DOC) mg/l (C), Chlorophyll-a μg/l (D) sampled at flooded floodplain agricultural wetlands, perennial channel, and Sacramento River habitats. Note that y-axis scales are variable between graphs so that differences in habitats may be distinguished.

Chlorophyll values were highest in the perennial drainage canal and lowest in the floodplain agricultural wetland (Fig 4). Chlorophyll declined throughout the study in the perennial drainage canal from a high of 19.2 μg/l to a low of 5.5 μg/l. Sacramento River chlororphyll values increased from the initial sampling 1.4 μg/l and slightly decreased prior to the final sampling event, to 3.7 μg/l. In the floodplain agricultural wetland, chlorophyll values remained relativley small compared to the other two locations with little change (0.2 to 0.7 μg/l).

### Zooplankton and phytoplankton biomass and vital rates

A total of 18, 20, and 15 taxa were identified from the floodplain agricultural wetland, perennial drainage canal and Sacramento River, respectively, throughout the study (Table 3). Mean zooplankton density for the three sample periods was 81,031 individuals/m^3^ in the floodplain agricultural wetlands, 11,831 individuals/m^3^ in the perennial channel, and 1,529 individuals/m^3^ in the Sacramento River. Numbers of zooplankton in the floodplain agricultural wetland declined from a high of 103,714 individuals/m^3^ to a low of 60,321 individuals/m^3^ throughout the study, yet remained much higher than either the perennial drainage canal or the Sacramento River habitats (Fig 5).

**Table 3.**
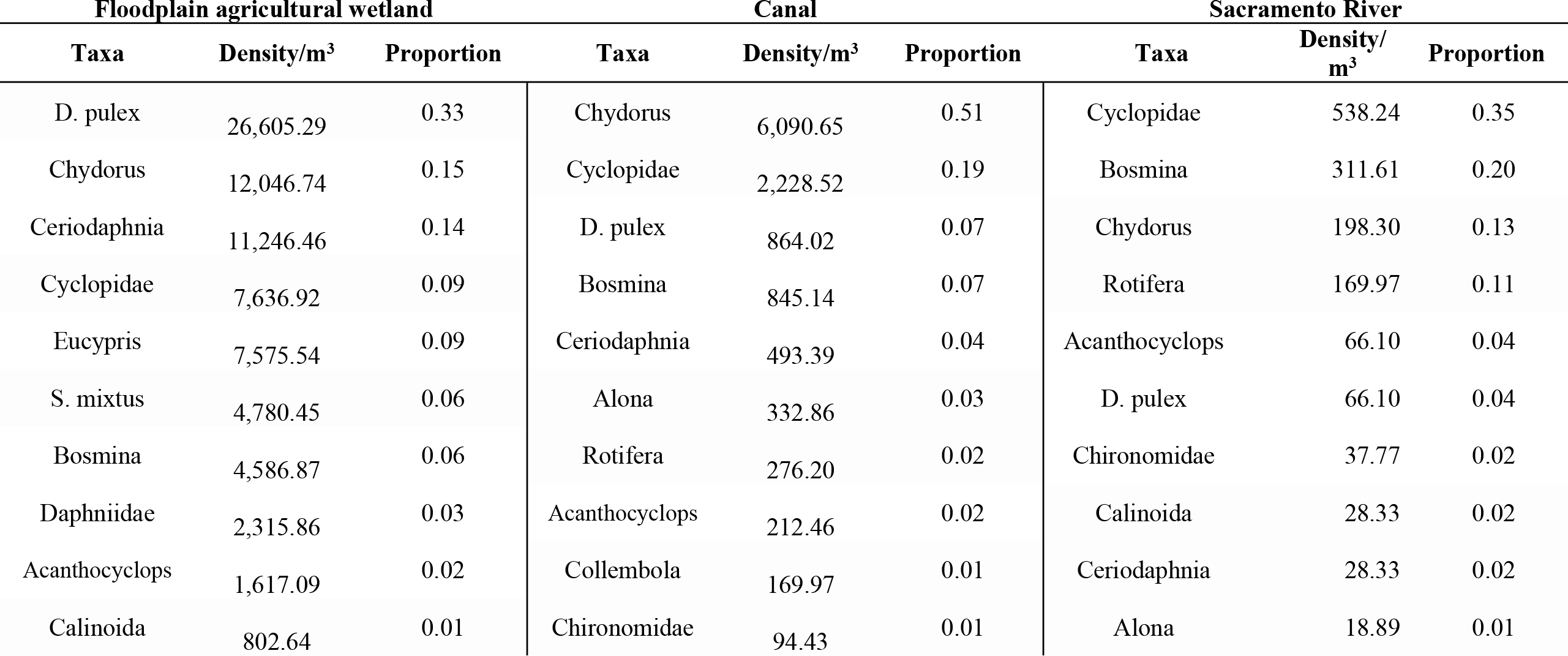
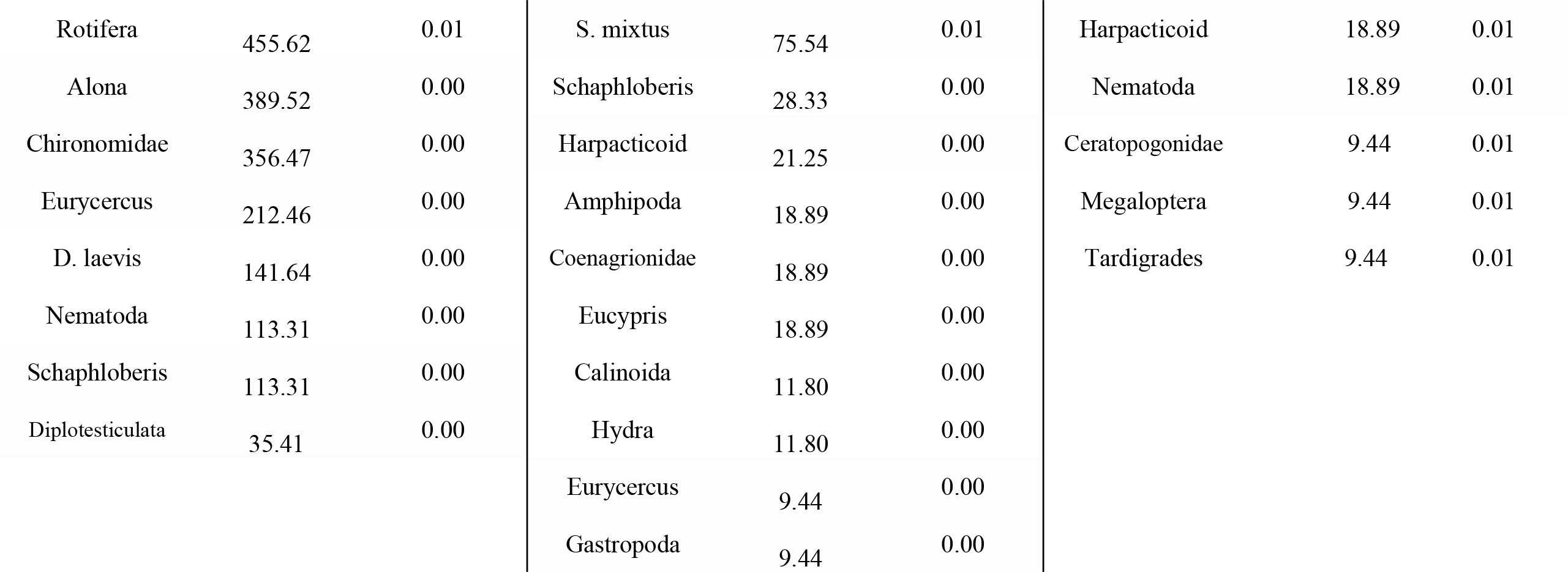
Taxa, mean density, and proportion of zooplankton collected during three sampling periods from the floodplain agricultural wetland, perennial channel, and Sacramento River habitats. Samples were collected adjacent to the fish enclosures.

**Fig 5.**
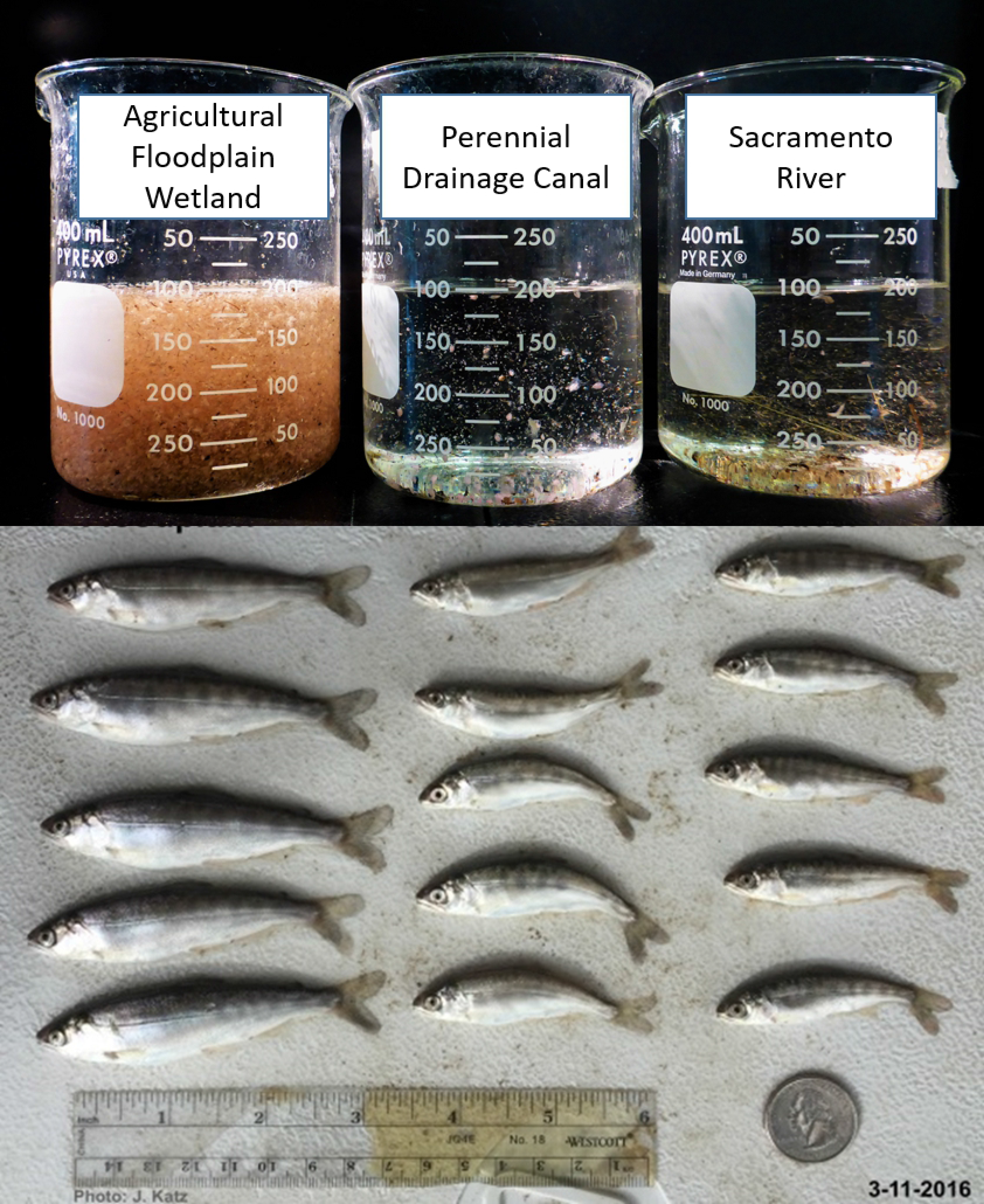
Estimated zooplankton density (individuals/m^3^) in the floodplain agricultural wetland, canal, and Sacramento River habitats.

Zooplankton grazing to primary production (ZG:PB) ratios varied over time and between habitats. The floodplain agricultural wetland ratio was the highest throughout the study but declined from a high of 6.67 to a low of 0.96 (Fig 6). Both the perennial drainage canal and Sacramento River locations remained well below a ratio of 1 indicating that there was more primary production than was estimated to be consumed by zooplankton.

**Fig 6.**
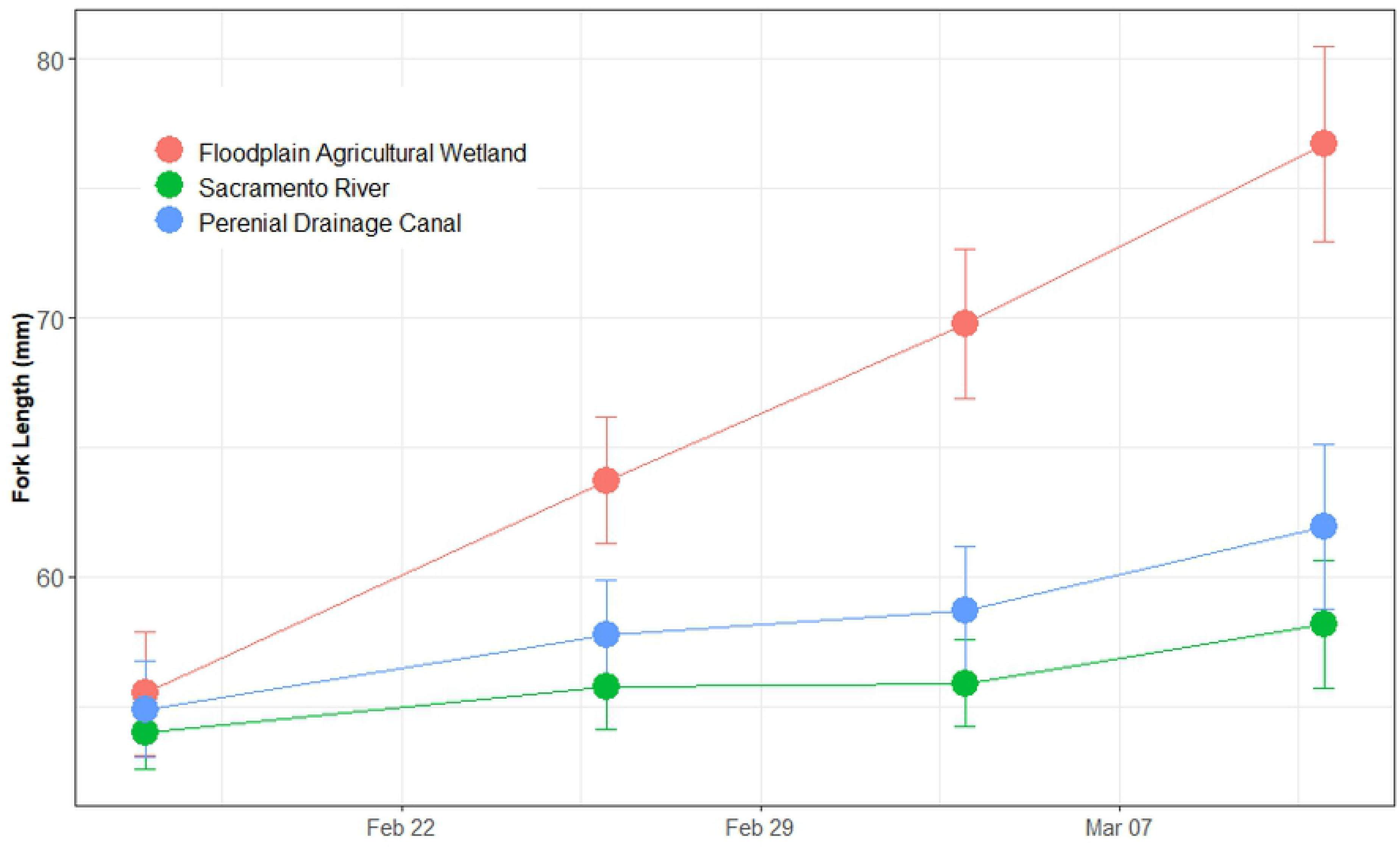
Ratio of zooplankton grazing: plankton biomass in floodplain agricultural wetlands, perennial channel, and Sacramento River. A value greater than 1 indicates there is more grazing than standing biomass of plankton, thus an alternate source of carbon than phytoplankton.

### Aquatic metabolism

Model results found that both the floodplain agricultural wetland and perennial drainage canal had similar daily average gross primary production values of 2.62 (+/− 1.59 sd) g/m^2^/d and 2.59 (+/− 1.53 sd) g/m^2^/d respectively while the Sacramento River was much lower 0.62 (+/− 0.79 sd) g/m^2^/d (Fig 7). Respiration values varied greatly between the three habitat types with the floodplain being the most negative at −10.17 (+/− 3.58 sd) g/m^2^/d (more respiration) and the Sacramento River site being the least negative −0.64 (+/− 0.98 sd) g/m^2^/d (less respiration). When respiration is subtracted from gross primary production net primary production (NPP) is the result. NPP values were estimated to be −7.56 (+/− 3.74 sd) g/m^2^/d, −3.27 (+/− 2.05 sd) g/m^2^/d, and −0.02 (+/− 1.66 sd) g/m^2^/d in the floodplain agricultural wetland, perennial drainage canal and Sacramento River respectively (Fig 7). Results from stream metabolism modeling had a difficult time with the variability in dissolved oxygen in the floodplain agricultural wetland. The modeled results failed to reach the minimum dissolved oxygen values observed in the agricultural wetlands. This likely resulted in an underestimation of the respiration and net ecosystem productivity output values (Fig 8).

**Fig 7.**
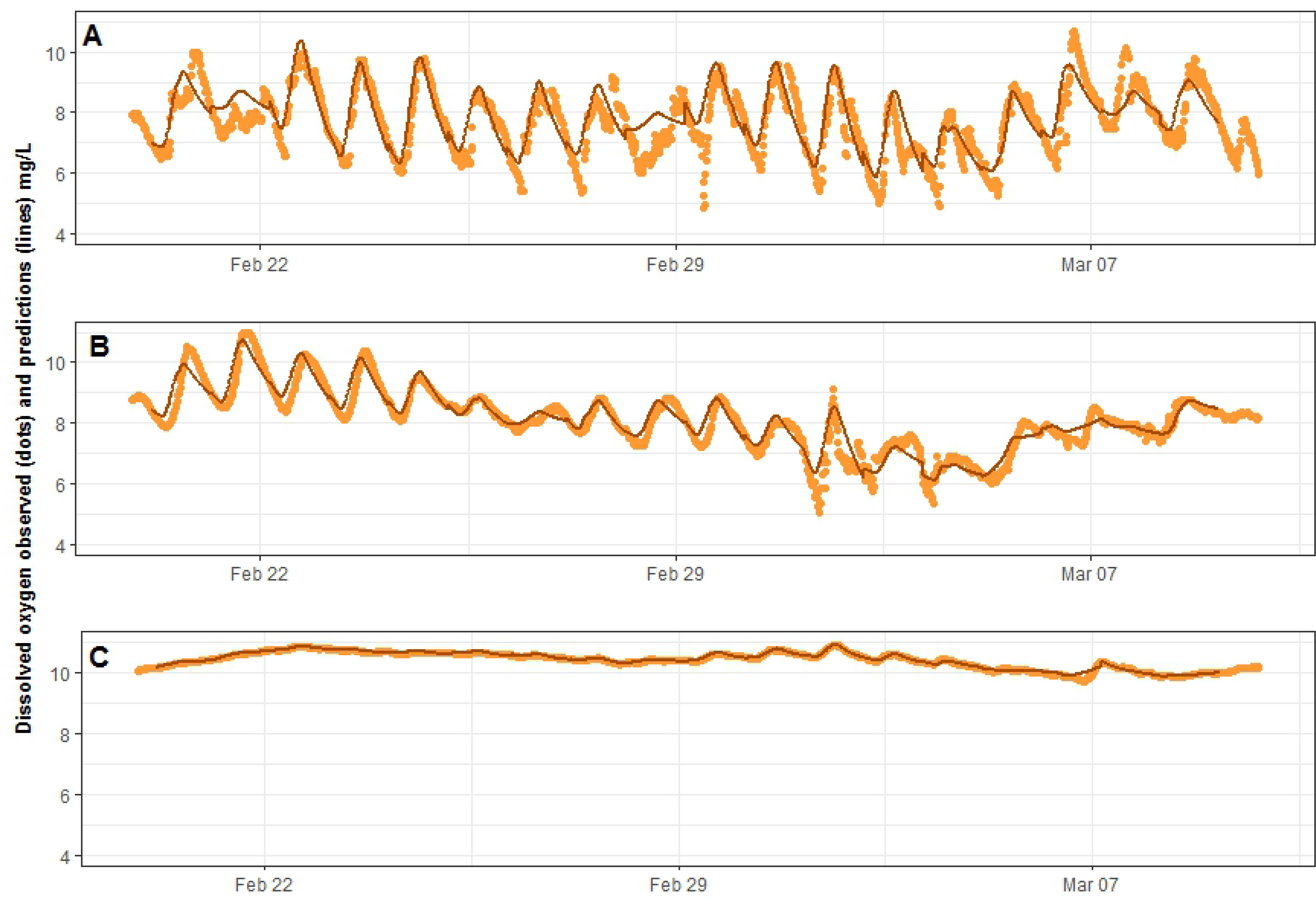
Model output of A) gross primary productivity (GPP, gC/m^2^/d), system respiration, net ecosystem productivity (NEP, gC/m^2^/d), and k-constant (m/d).

**Fig 8.**
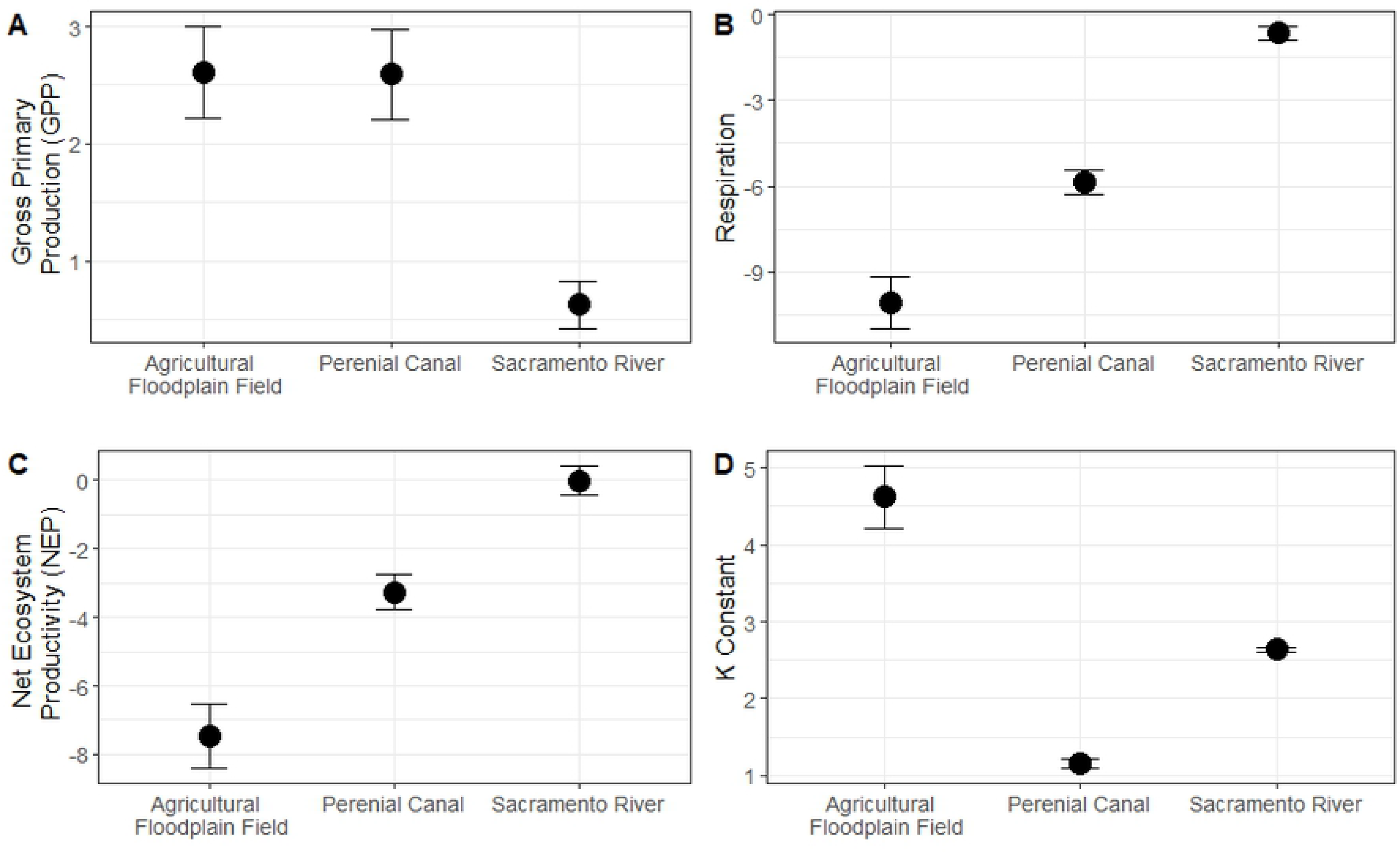
Actual and modeled Dissolved Oxygen data for a) floodplain agricultural wetland, b) perennial channel, and c) Sacramento River output from Streammetabolizer package in R.

### Fish growth

In the floodplain agricultural wetland habitat, six fish escaped through a hole in one of the enclosures after initial stocking, and one fish escaped and one fish died prior to the final sampling event, leaving a total of 22 fish measured through the entire study. Of the seven escaped fish, five were later recaptured following the study during the draining of the agricultural wetland. In the Sacramento River habitat, one fish died during removal from the enclosure for measurement, for a total of 29 fish that were measured through the studies duration. All 30 fish initially planted in the perennial drainage canal habitat were present on the final day of the study. Juvenile Chinook Salmon averaged 54.8 mm (+/− 2.00 sd) across all habitats when initially placed into the enclosures (Fig 9). Throughout the study, fish in the floodplain agricultural wetland, perennial channel, and Sacramento River grew in fork length, on average, 0.93 (+/−0.15 sd) mm/day, 0.31 (+/− 0.10 sd) mm/day, 0.18 (+/− 0.09 sd) mm/day, respectively. At the conclusion of the study, the average fork lengths were floodplain agricultural wetland = 76.7 mm (+/− 3.78 sd), perennial drainage canal = 62.0 mm (+/− 3.17 sd), and Sacramento River = 58.2 mm (+/− 2.45 sd) (Fig 9). Fish in the floodplain agricultural wetlands were significantly longer in fork length than the fish in either the perennial drainage canal or the Sacramento River (p < 0.0001) and the perennial drainage canal fish were significantly longer in fork length that the fish in the Sacramento River (p<0.0001) (Fig 10).

**Fig 9.**
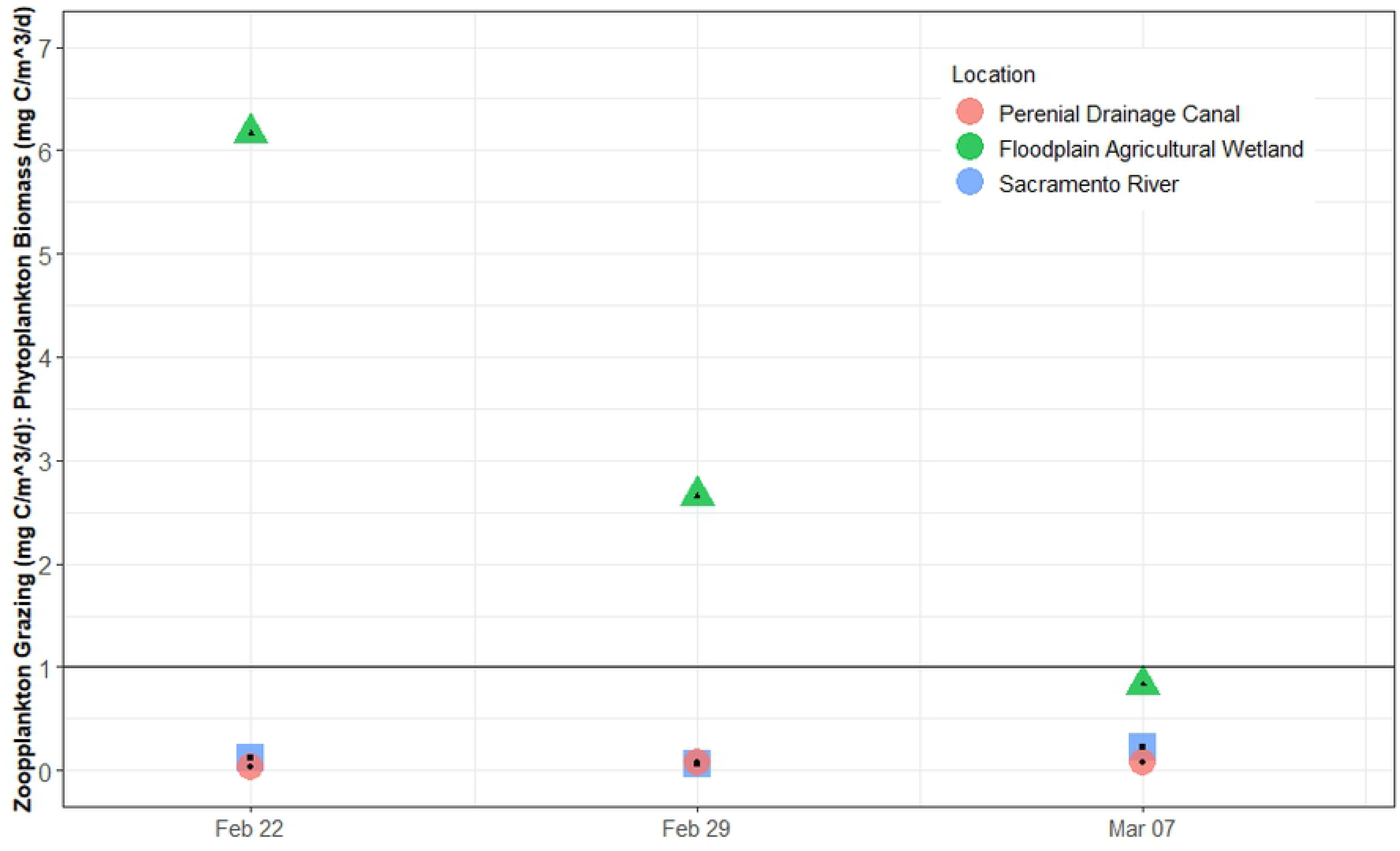
Fork length of juvenile fish reared in a flooded floodplain agricultural wetland, perennial drainage channel, and Sacramento River. Circles are habitat mean fork length, and error bars are standard deviation.

**Fig 10.**
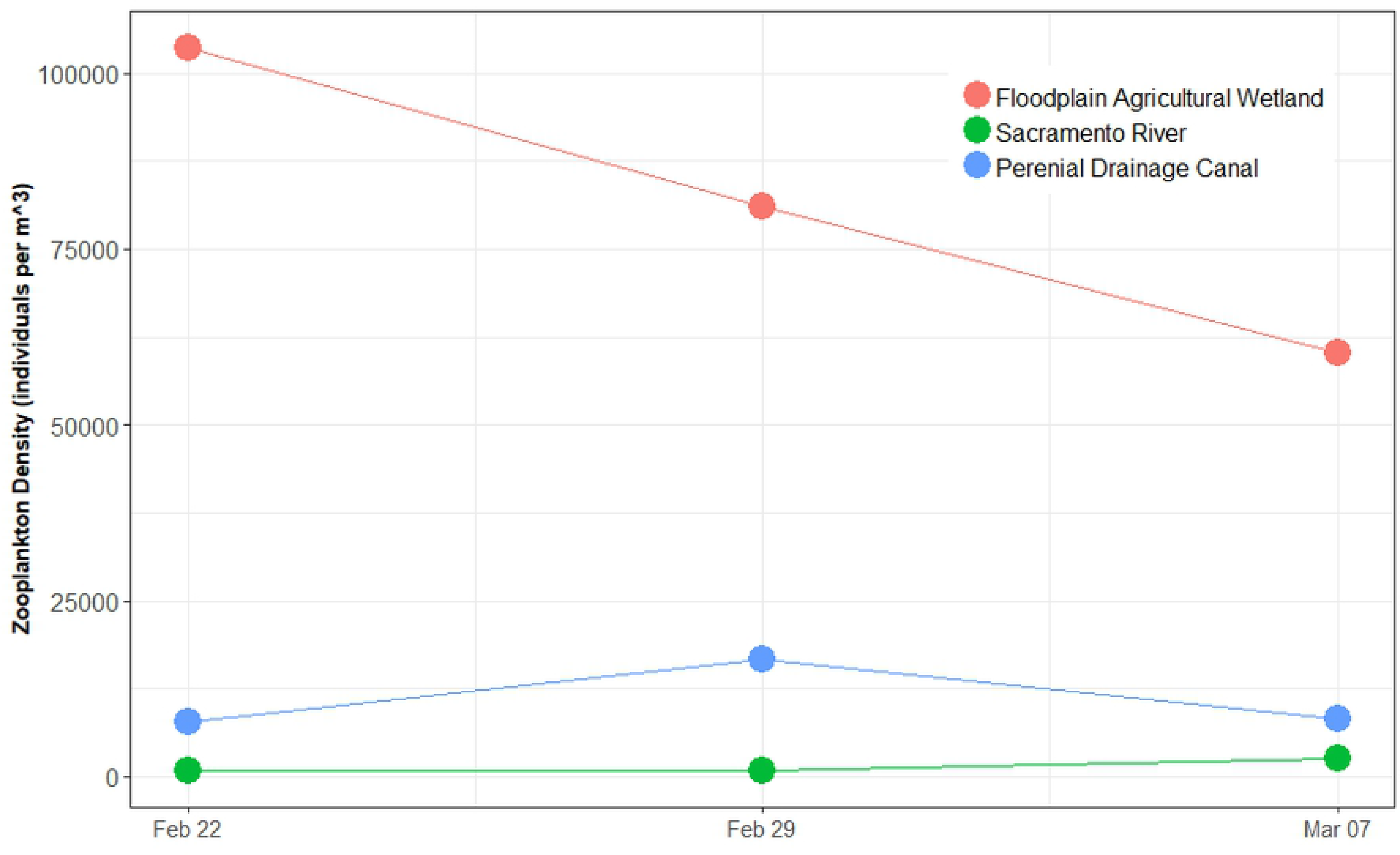
Photo of zooplankton sample and representative fish from each of the three habitat types, flooded agricultural wetland, perennial drainage channel, and Sacramento River. Note abundance of zooplankton and associated size of fish from the flooded agricultural floodplain habitat compared to the two other locations.

## Discussion

The purpose of our study was to examine food web responses across a range of habitat conditions in a river-floodplain complex. Like many large-river systems, the Sacramento River and its adjacent floodplain (Yolo Bypass) have been heavily modified for flood management and agriculture [6]. For example, levees and weirs have substantially reduced the connectivity between the Sacramento River and the Yolo Bypass, reducing the frequency and duration of seasonal inundation. Nonetheless, our study suggests that even heavily managed floodplains such as Yolo Bypass retain habitat attributes that enhance the productivity and diversity of the food web. Overall, we found support for our hypotheses that off-channel agricultural wetland habitat generates higher densities of zooplankton and increased growth rates of juvenile salmon as compared to adjacent perennial canal and river channels. While our study design relied on the use of caged hatchery fish and a heavily managed agricultural wetland as part of the habitat comparisons, these results are consistent with observations of zooplankton densities and wild fish growth rates during more natural uncontrolled flood events [17, 31, 32]. Below we describe some of the habitat attributes which may be responsible for these differences.

Primary production is generally regulated by either bottom up limitations (nutrients) or top down grazing pressure from primary consumers, in this case zooplankton [33, 34]. The floodplain habitat contained high densities of zooplankton yet exhibited low concentrations of chlorophyll. While higher zooplankton levels on floodplain habitat than adjacent channels is consistent with earlier studies [17, 32], the lower chlorophyll a levels in the agricultural wetland is different than observations during flood events [32]. If primary production rates are high, intense grazing pressure from abundant zooplankton can keep observable chlorophyll biomass low. However, this is not likely to have occurred here as nitrate-nitrogen values were also low (0.00-0.01 mg/L) limiting rates of primary production to far below what would be needed to sustain the abundance of zooplankton observed. Additionally, modeled oxygen consumption rates were observed to be as much as six times primary production rates (Fig 8) suggesting that respiration in the system far exceeded oxygen production from photosynthesis. Thus, the highly productive floodplain food web is not likely being fueled by autotrophic production. This leads us to conclude that a detrital-based heterotrophic food web is strongly contributing to production of zooplankton densities approximately 53 times as dense, on average, than those produced by the in-channel food web of the adjacent Sacramento River. Ephemerally abundant floodplain food resources may be utilized by migratory fish that gain access to hydrologically activated floodplain habitats. Floodplain-derived food web resources may also be exported to downstream water bodies as floodplains drain, thereby subsidizing in-river food webs and making floodplain-derived food web resources available to fish populations confined to downstream river channels.

Differences in basal carbon resources were likely the largest contributing factors to observed differences in zooplankton abundance and juvenile salmon growth between the habitats, with temperature and residence time of water also playing contributing roles. During this study, in the Sacramento River the primary source of carbon appears to be in situ primary production by autotrophic phytoplankton. Both autotrophic and heterotrophic pathways appear to be important in the perennial channel, while floodplain carbon appears to be primarily detrital.

Inundation of floodplains facilitates the decomposition of terrestrial vegetation and allows soils to leach labile carbon into the water column. These detrital carbon sources as well as methane produced in the oxygen-poor wetland habitats and support single celled organisms and methane oxidizing bacteria [11]. Heterotrophic pathways can fuel a highly productive food web even in the absence of high phytoplankton production [13]. In off-channel shallow lakes in the Amazon River, up to 84% of the amino acids in fish were derived from methane oxidizing bacteria [11]. Previous work on the Yolo Bypass has found that a detritivorous chironomid midge was the primary food source for juvenile salmonids and other fishes utilizing the flooded Yolo Bypass [16, 17].

Longer residence times of water on the floodplain also likely contributed to higher zooplankton densities by increasing the labile carbon availability for microbial decomposition. Water temperatures in the shallowly inundated floodplain agricultural wetland and the perennial drainage canal were warmer and more variable compared to those in the Sacramento River (Fig 2), which is consistent with earlier studies in this region [17]. When water temperatures are higher, yet within physiological tolerances, and food resources are abundant, juvenile salmon growth rates can exceed those of fish in cooler habitats [35]. Warmer temperatures also allow for increase microbial production, thus more basal food web production [36]. While water temperatures were similar between the floodplain agricultural wetland and perennial channel, zooplankton densities were much greater on the floodplain, resulting in elevated foraging efficiencies and higher growth rates observed in the juvenile salmon (Fig 9).

Chlorophyll is often used a proxy for productivity in aquatic ecosystem. This may be partially due to a widely-held conceptual model that focuses on autotrophic production as the base of aquatic food webs. The central role of chlorophyll as the primary measure of aquatic ecosystem productivity may be further entrenched by the relative ease of measuring chlorophyll while obtaining a direct measure of the heterotrophic production is inherently difficult. Our results suggest that heterotrophic production is the primary base of the food web in the floodplain food web studied here. These results also make clear that different habitats, even those that are relatively close together and intermittently hydrologically connected, can support dramatically different food webs and ultimately provide very different growth potential for juvenile salmon. The relative importance of floodplain-derived food resources in the salmonid life-cycle suggests that detrital food webs may deserve greater scrutiny as important engines for the production aquatic biomass, especially fish. Fish in the inundated floodplain agricultural wetland grew significantly faster (5x) than fish in either the perennial drainage canal or Sacramento River (Fig 9). Food resources in the Sacramento River were generally sparse compared to those on the floodplain corroborating past studies that have shown fish rearing in the river to have slower growth rates when compared to those rearing in adjacent off-channel habitats [18, 37].

Hydrologic conditions typifying the ephemeral floodplain–shallower depths, warmer water, longer residence times and detrital carbon sources compared to deeper, colder, swifter water and an algal-based carbon source in the adjacent river channel–appear to facilitate the dramatically higher rates of food web production observed in floodplain verses river channel habitats. These results suggest that hydrologic patterns associated with winter flooding provide Mediterranean river systems access to detrital carbon sources that may be important energy sources for the production of fisheries resources and biomass. Future conservation, farm, water and fisheries management actions informed by this study can be tailored to the production of aquatic food resources to benefit fisheries and imperiled native fish species.

## Acknowledgements

We would like the thank John Brennan, David Katz, Huey Johnson, and Laney Thorton of Cal Marsh and Farms whose vision make this work possible. We would also like to thank Conaway Ranch for access for the study. Funding was provided by the California Department of Water Resources, the Water Foundation, and California Trout. The US Bureau of Reclamation funded previous year’s research that made this work possible. We would also like to thank Miranda Tilcock and Rosa Cox for fieldwork, sample identification and data collection and Randy Dahlgren for water quality assistance and general guidance throughout the project.

